# Evaluating denoising strategies in resting-state fMRI in traumatic brain injury (EpiBioS4Rx)

**DOI:** 10.1101/2021.12.10.472139

**Authors:** Marina Weiler, Raphael F. Casseb, Brunno M. de Campos, Julia S. Crone, Evan S. Lutkenhoff, Paul M. Vespa, Martin M. Monti, for the EpiBioS4Rx Study Group

**Author notes:** Co-first authors contributed equally (Marina Weiler, Raphael F. Casseb). Correspondence to: Martin M. Monti, Department of Psychology, University of California Los Angeles, Los Angeles, CA 90095, USA.

## Abstract

**Objective:** Resting-state functional MRI is increasingly used in the clinical setting and is now included in some diagnostic guidelines for severe brain injury patients. However, to ensure high-quality data, one should mitigate fMRI-related noise typical of this population. Therefore, we aimed to evaluate the ability of different preprocessing strategies to mitigate noise-related signal (*i*.*e*., in-scanner movement and physiological noise) in functional connectivity of traumatic brain injury patients.

**Methods:** We applied nine commonly used denoising strategies, combined into 17 pipelines, to 88 traumatic brain injury patients from the Epilepsy Bioinformatics Study for Anti-epileptogenic Therapy clinical trial (EpiBioS4Rx). Pipelines were evaluated by three quality control metrics across three exclusion regimes based on the participant’s head movement profile.

**Results:** While no pipeline eliminated noise effects on functional connectivity, some pipelines exhibited relatively high effectiveness depending on the exclusion regime. Once high-motion participants were excluded, the choice of denoising pipeline becomes secondary - although this strategy leads to substantial data loss. Pipelines combining spike regression with physiological regressors were the best performers, whereas pipelines that used automated data driven methods performed comparatively worse.

**Conclusion:** In this study, we report the first large-scale evaluation of denoising pipelines aimed at reducing noise-related functional connectivity in a clinical population known to be highly susceptible to in-scanner motion and significant anatomical abnormalities. If resting-state functional magnetic resonance is to be a successful clinical technique, it is crucial that procedures mitigating the effect of noise be systematically evaluated in the most challenging populations, such as traumatic brain injury datasets.

## 1. INTRODUCTION

Over the last few decades, the assessment of spontaneous oscillations in the blood oxygenation level-dependent (BOLD) measured by resting-state functional magnetic resonance (rsfMRI) has increasingly been used to aid diagnosis and prognosis in neurological disorders (Baker et al., 2014; de Vos et al., 2018; Franzmeier et al., 2020; Wolters et al., 2019; Woodward et al., 2012). Yet, despite the appeal and wide adoption of this technique, it suffers from significant limitations for distinguishing oscillations associated with neural activity from those induced by non-neural sources (Birn, 2012; Murphy et al., 2013; Power et al., 2017). In-scanner head motion can systematically generate artifactual correlations across brain regions and spurious functional connectivity (FC) results regardless of how they are assessed (e.g., seed-based analysis, graph theory, amplitude of low frequency fluctuations)(Power et al., 2012; Satterthwaite et al., 2012; Van Dijk et al., 2012).

In the context of severe brain injury and disorders of consciousness, some international guidelines (Kondziella et al., 2020) now suggest incorporating rsfMRI in the diagnostic process given its ability to complement bedside neurobehavioral assessments and provide prognostic information (Demertzi et al., 2019; Madhavan et al., 2019; Silva et al., 2015; Vanhaudenhuyse et al., 2010). However, this patient group is well known to exhibit high incidence of in-scanner motion during data acquisitions (Hannawi et al., 2016; Monti et al., 2015), which can corrupt estimates of FC. While this issue could be mitigated with the use of sedative agents, these will affect any subsequent analysis of brain network function (Monti et al., 2013), thus making the development of analytical approaches to mitigating in-scanner motion a more desirable strategy. In this sense, a large number of analysis pipelines have been proposed to address the issue (Muschelli et al., 2014; Power et al., 2015). However, most of this work has been developed and evaluated in neurotypical individuals or clinical populations that do not usually present significant anatomical abnormalities (Burgess et al., 2016; Ciric et al., 2018; Parkes et al., 2018; Power et al., 2020; Raval et al., 2020). No pipeline has ever been validated with respect to patients exhibiting the degree of in-scanner motion (Hannawi et al., 2016; Monti et al., 2015) and the extensive brain pathology (such as atrophy and trauma-induced deformations) known to lead to sub-optimal and biased performance of conventional analysis software (Lutkenhoff et al., 2014). If rsfMRI is to be a successful technique used in routine clinical practice (Kondziella et al., 2020), it is crucial that procedures mitigating the effect of noise be systematically evaluated also in the most challenging populations.

To address this gap, we extend a prior large-scale evaluation of different pipelines (Parkes et al., 2018) to the very challenging population of moderate-to-severe traumatic brain injury (TBI) to provide a quantitative comparative assessment of different denoising strategies. Specifically, we applied nine commonly used denoising strategies, combined into 17 pipelines, to TBI patients from the Epilepsy Bioinformatics Study for Anti-epileptogenic Therapy clinical trial (EpiBioS4Rx) (Vespa et al., 2019) and evaluated the ability of each one to remove noise from the BOLD signal. We conclude by providing a framework for clinicians and translational scientists interested in using fMRI to select the pipeline that balances the ability to mitigate noise with the constraints and aims of their study.

## 2. METHODS

### 2.1 Subjects

This study included 88 patients from the EpiBioS4Rx dataset, a longitudinal study that aims to discover and validate observational biomarkers of epileptogenesis after TBI (Vespa et al., 2019). As described elsewhere, patients were enrolled across 12 sites within 72 hours following TBI involving frontal and/or temporal hemorrhagic contusion, according to criteria previously published (Vespa et al., 2019). Our sample consisted of 21 females and 67 males, with mean age 41.1 (7-84) years, level of consciousness after TBI measured by the Glasgow Coma Scale (Teasdale and Jennett, 1974) 7.8 (1-15), and time since injury 11 (0-36) days. Informed consent was obtained from a surrogate family member or legally authorized representative, using IRB-approved consent methods.

### 2.2 Image Acquisition and Processing

Data were acquired on 1.5 or 3T MR system, including an anatomical (T1-weighted) and functional (T2*-weighted echo planar images) acquisitions (See **Suppl. Tables 1** and **2** for detailed parameter listing.) Data were processed using code adapted from (Parkes et al., 2018) (https://github.com/lindenmp/rs-fMRI). Before temporal and spatial filtering, preprocessed data were submitted to denoising strategies (see below).

### 2.3 Denoising Strategies

Denoising is achieved by removing variance attributable to head and respiration/cardiac -induced motion from the BOLD signal. What is debated is how to best measure, operationalize, and remove these sources of noise. **Table 1** summarizes the denoising approaches used in our analysis. We combined these approaches into 17 pipelines, as done in prior work that used a different clinical sample (Parkes et al., 2018).

**Table 1:** Denoising strategies.

|  |  |
| --- | --- |
| **Head displacement** | **Head Motion Parameters (HMP):** Six parameters (3 rotations and 3 translations about/along the X-, Y-, and Z-axes) included as noise regressors (6HMP)<sup>36</sup>. Additional regressors derived from the six parameters (*e.g.*, temporal and quadratic terms of each parameter, as well as their difference) are often included to account for delayed and nonlinear motion-induced spin history effects (24HMP)<sup>31</sup>.<br>**Spike Regression**<sup>31</sup>: For each volume containing excessive motion, a separate regressor is generated containing a value of 1 at that volume, and 0 at all others. Volumes are considered contaminated if $FD_{Jenk}^{28} > 0.25$ mm. $FD_{Jenk}$ represents the root mean squared of the 6 motion parameters<sup>28</sup>.<br>**Scrubbing**<sup>29</sup>: Each volume containing excessive motion is removed from the time-series if $FD_{Power}^9 > 0.2$ mm or $DVARs^{9,27} > 2$ %. After removal, uncontaminated segments of BOLD data lasting fewer than 5 contiguous volumes are also removed. $FD_{Power}$ represents the sum of the absolute values of the differentiated realignment estimates (by backward differences) at every time-point<sup>9</sup>. |
| **Physiology-related** | **Physiological Regressors (2phys):** Regression of the average signal from WM and CSF, tissues not expected to exhibit BOLD oscillations tied to neural activity.<br>**Anatomical Component Based Correction (aCompCor<sup>37</sup>/aCompCor50<sup>38</sup>):** This approach involves extracting orthogonal components of temporal variance from voxel-wise time series for the WM and CSF masks separately. Then, either the five components with greater eigenvalue for each tissue are included in the denoising regression (aCompCor), or as many components as needed to cumulatively explain at least 50% of the variance in each tissue (aCompCor50). |
| **Mixed approaches** | **Global Signal Regression (GSR):** Regression of the average signal across all the voxels of the brain.<br>**ICA-based Automatic Removal Of Motion Artifacts (ICA-AROMA<sup>39</sup>):** Automated data-driven method to identify and remove via regression motion-related independent components. |

Abbreviations: FD = framewise displacement; DVARs = derivative of the root mean squares variance over voxels; BOLD = blood oxygenation level-dependent; WM = white matter; CSF = cerebrospinal fluid; ICA = independent component analysis.

### 2.4 Head Movement Estimation (in-scanner motion)

As shown in **Table 1**, some pipelines rely on the ability to pinpoint volumes corrupted by excessive motion. In general, motion in a volume is quantified by the Derivative of the root mean squares VARiance over voxelS (DVARS) (Power et al., 2012; Smyser et al., 2011) and Framewise Displacement (FD) (Jenkinson et al., 2002; Power et al., 2012), which were used here for spike regression and scrubbing approaches. In addition, we calculated in-scanner head movement for each patient, used to calculate quality control measures, and classify each patient under an exclusion regime. For that, we used the mean FD_Jenk_ (Jenkinson et al., 2002) across all volumes (hereafter, mFD).

### 2.5 Quality Control (QC) Measures

After image preprocessing, we used a template containing 333 cortical regions (ROIs) (Gordon et al., 2016) to define the areas to extract gray matter (GM)-weighed denoised time-series for further analysis. We then calculated FC as Pearson’s correlation coefficient between each pair of ROI time-series, then implemented a Fisher’s r-to-z transformation. The FC matrices obtained following each denoising pipeline were then used to evaluate the ability of each pipeline to remove noise-induced correlations by the two quality control measures described below.

#### 2.5.1 QC-FC correlation

Represents the correlation between FC and in-scanner head motion (mFD) since non-neuronal fluctuations can increase the apparent FC between regions by introducing spurious common variance across time series. Here, we calculated Pearson’s correlation coefficient between each pair of ROIs FC and the mFD across patients.

#### 2.5.2 QC-FC distance-dependence

Indicates whether the correlation between FC and in-scanner head motion (mFD) is spatially structured – a known feature of motion-induced artifacts (Power et al., 2012; Power et al., 2014; Satterthwaite et al., 2012; Van Dijk et al., 2012). Here, we calculated the distance between ROIs as the Euclidean distance between the stereotaxic coordinates of the volumetric centers of ROI pairs. We quantified the relationship between this distance and the QC-FC correlation for each edge using Spearman’s rank correlation coefficient (ρ) due to the non-linearity of some associations.

In addition, we measured the ability of each pipeline to retain statistical power during the denoising process:

#### 2.5.3 loss of temporal degrees of freedom (tDOF-loss)

Represents the amount of tDOF lost due to the removal of time points and/or to the number of regressors used to denoise the data (**Suppl. Table 3**).

### 2.6 Participant Exclusion Regimes

Finally, it is debated how to determine the threshold at which a subject contains excessive motion and thus should be discarded from any analysis. We compared the performance of all pipelines under three different participant exclusion regimes: (*i*) censoring-based, (*ii*) lenient, and (*iii*) stringent (**Table 2** shows criteria for subject exclusion in each regime).

**Table 2:** Participant exclusion regimes and their criteria for exclusion:

| Regime | Exclusion criteria |
| --- | --- |
| **Censoring-based**(Satterthwaite et al., 2013; Van Dijk et al., 2012) | Excluded subjects when less than 4 minutes of non-contaminated volumes remained after volume censoring (< 4 minutes of data). |
| **Lenient**(Satterthwaite et al., 2012) | Excluded subjects if:<br>(i) < 4 minutes of data; or<br>(ii) high levels of head gross motion, defined as mFD > 0.55mm. |
| **Stringent**(Satterthwaite et al., 2013) | Excluded subjects if:<br>(i) < 4 minutes of data;<br>(ii) mFD > 0.25 mm;<br>(iii) more than 20% of the volumes presented $FD_{Jenk} > 0.2$ mm; or<br>(iv) any volume presented $FD_{Jenk} > 5$mm. |

Abbreviations: FD = framewise displacement; mFD = mean framewise displacement.

## 3. RESULTS

### 3.1 Head Movement and Participant Exclusion

As shown in **Fig. 1a**, censoring-based, lenient, and stringent regimes resulted in the exclusion of 8, 11, and 32 patients (9 %, 12.5 %, and 36 %, respectively). As expected, participants under the stringent regime presented significantly smaller mFD compared to censoring-based and lenient regimes (**Fig. 1b**, Kruskal-Wallis test, *H(2)* = 9.791, *p* = 0.0075; mean rank mFD 116.45 for censoring-based, 113.18 for lenient and 85.00 for stringent. Pairwise comparisons Bonferroni-adjusted for multiplicity: stringent vs lenient, *p* = 0.028; stringent vs censoring-based, *p* = 0.01; lenient vs censoring-based, *p* = 1). In-scanner head movement did not correlate with Glasgow Coma Scale (Pearson *r(84)* = 0.080, *p* = 0.462), age (Pearson *r(85)* = 0.139, *p* = 0.201), nor time since injury (Pearson *r(79)* = 0.128, *p* = 0.253). A logistic regression was performed to ascertain the effects of age, gender, time since injury, and Glasgow Coma Scale on the likelihood that participants would fall into the Stringent regime. The logistic regression model was not statistically significant, χ2(4) = 6.051, *p* = .195. The model explained 10% (Nagelkerke *R*^*2*^) of the variance in Stringent regime and correctly classified 63% of cases.

**Fig 1:**
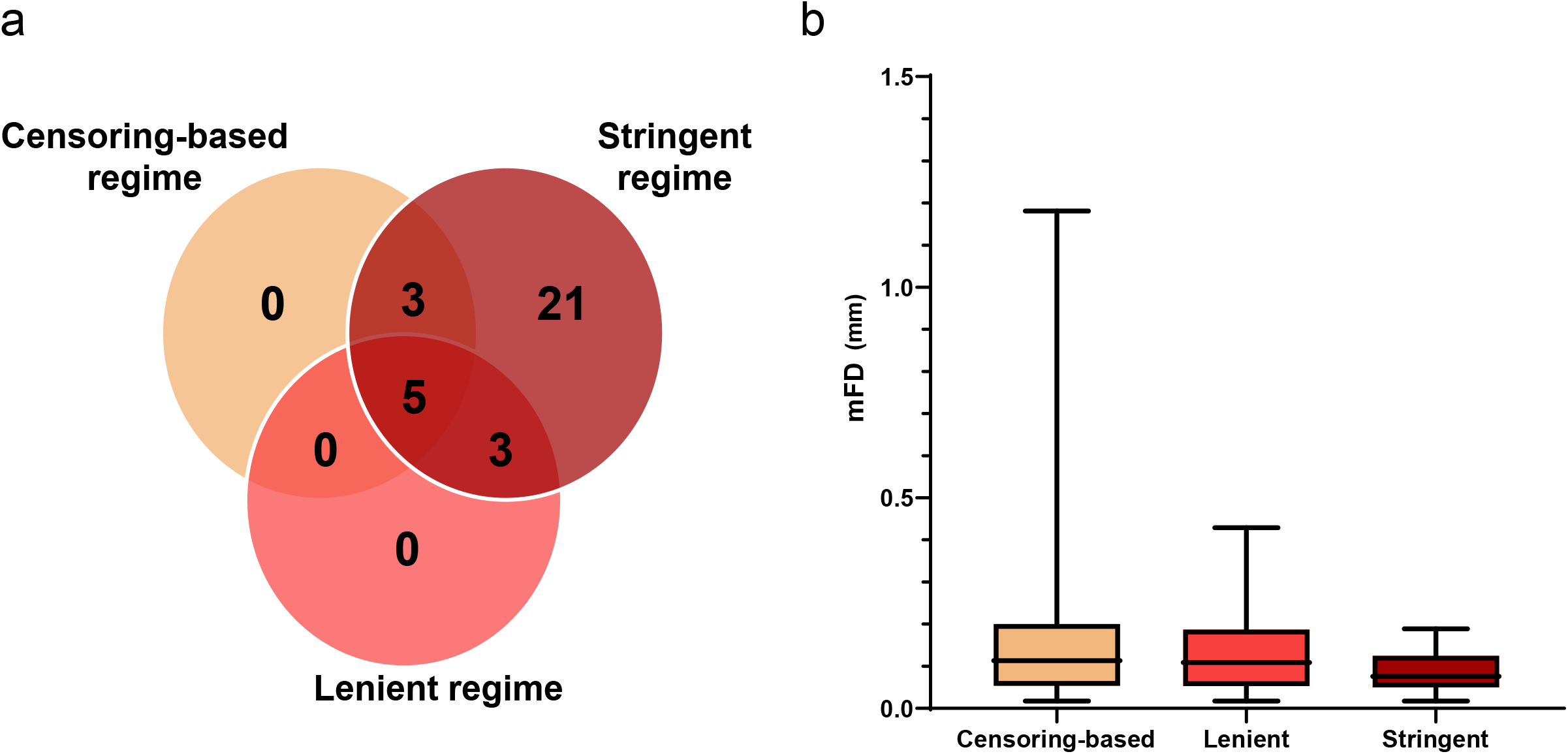
(a) Number of participants excluded in each regime; (b) Box plots of the mFD values for each regime.

### 3.2 Quality Control Measures

As shown in **Fig. 2** and **3**, consistent with prior work (Parkes et al., 2018), while no pipeline entirely eliminated noise-related effects on FC patterns, some pipelines exhibited relatively high effectiveness at mitigating it depending on the exclusion regime.

**Fig 2:**
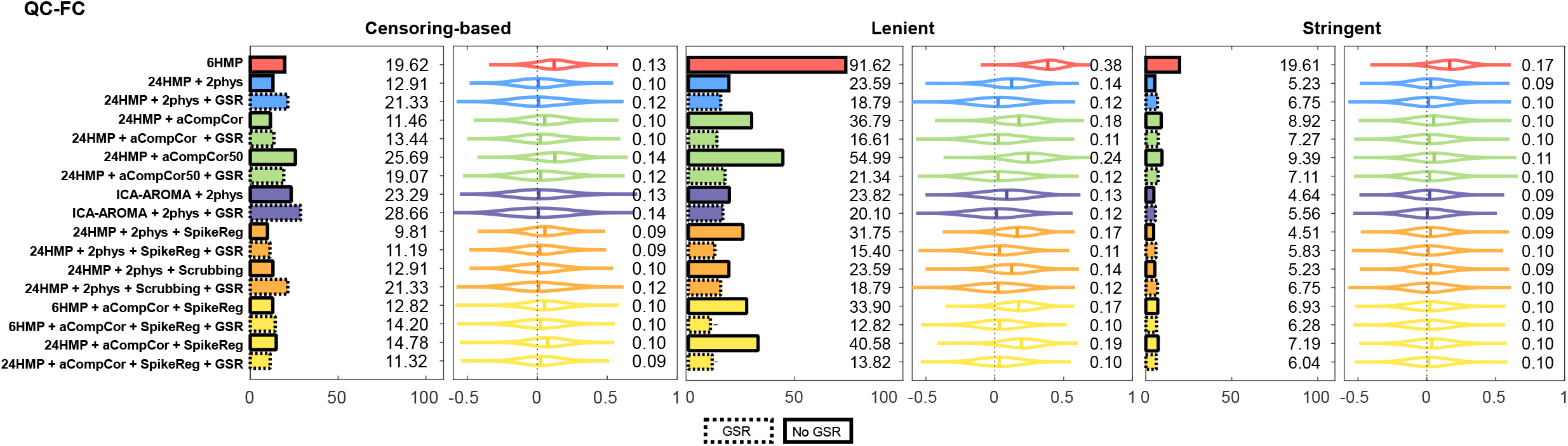
QC-FC correlations under the three regimes of participant exclusion. On the left of each panel, results are shown as the proportion of significant FCs that correlated with the patient’s head movement (mFD), *p* < 0.05. On the right of each panel, results are shown as the full distribution of QC-FC, and the corresponding median value. Better denoising pipelines result in fewer correlations between FC and head movement, giving values closer to 0.

#### 3.2.1 QC-FC

Overall, no pipeline in any exclusion regime reduced the effect of noise to zero (**Fig. 2**). Nonetheless, our results show that some pipelines, under a given regime, perform better than others.

First, the censoring-based and lenient regimes resulted in approximately 10-29 % and 13-55 % proportion of significant correlations and absolute *r*-values between 0.09-0.14and 0.10-0.24, respectively. 6HMP pipeline was an outlier with ∼92 % proportion of QC-FC significant correlations and an absolute correlation between motion and FC of 0.38 in the lenient regime. In comparison, the stringent criterion resulted in lower QF-FC across all pipelines, reducing the correlations significantly to less than 10 % and median *r*-value between motion and FC to 0.09-0.11 (with the sole exception of the 6HMP, with ∼20 % significant correlations).

Second, within each of the three exclusion regimes, different strategies exhibit different effectiveness at mitigating noise. Overall, in the (*i*) censoring-based regime, the best performance was obtained with different combinations of 24HMP, aCompCor, and spike regression. Conversely, the three worst performing pipelines all featured data-driven methods, including aCompCor50 and ICA-AROMA. The addition of GSR generally resulted in the worsening of pipeline performance. Under the (*ii*) lenient regime, a very different pattern of results was observed. Overall, the best performing pipelines under this regime were the two featuring aCompCor, spike regression, and GSR, with either 6 or 24 HMP. At the opposite end of performance, pipelines without GSR underperformed those with GSR, and the pipeline with 6HMP alone resulted in the slightest mitigation of noise-induced effects on FC. The inclusion of GSR improved pipeline performance for all pipelines, with the greatest benefit observed for the aCompCor/aCompCor50 pipelines. Overall, in the (*iii*) stringent regime, the pipelines performed similarly one to another (with the sole exception of the 6HMP), and all pipelines performed better under the stringent regime than censoring-based and lenient regimes, reducing significantly QC-FC correlations. The inclusion of GSR barely changed any pipeline performance under this regime.

#### 3.2.2 QC-FC Distance-Dependence

As shown in **Fig. 3**, the proportion of statistically significant correlations between QC-FC and ROI distance for each denoising pipeline in each regime is comparable to prior validations (Parkes et al., 2018). Like QC-FC, the stringent regime reduced distance dependence on QC-FC the most (with an absolute average correlation of 0.06), followed by the lenient and the censoring-based criteria (absolute average correlation of 0.16 and 0.25, respectively).

**Fig 3:**
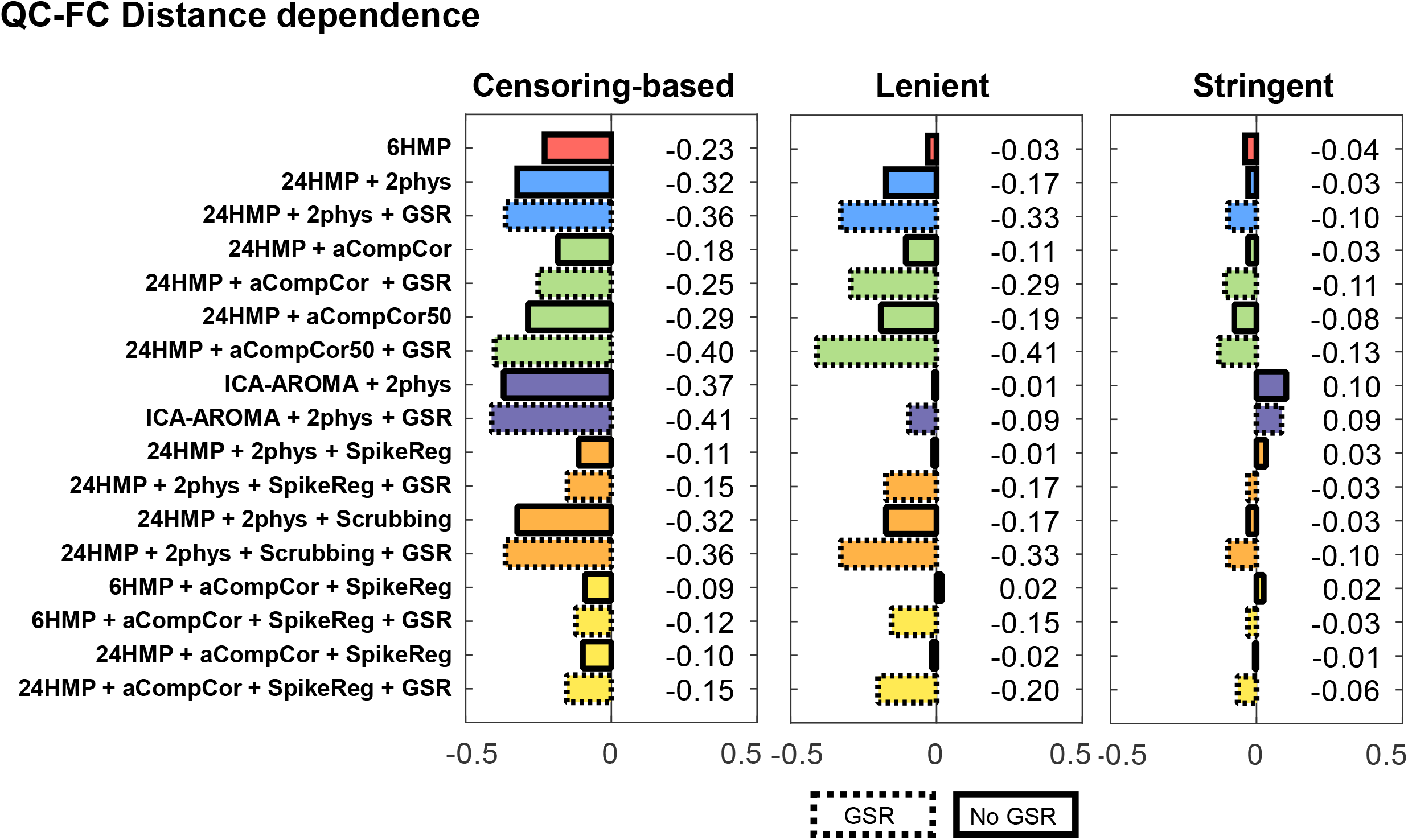
QC-FC distance-dependence under the three participant exclusion regimes. Results are presented as Spearman’s ρ correlation coefficient. Better denoising pipelines result in fewer correlations between FC and head movement, giving values closer to 0.

Specifically, in the (*i*) censoring-based regime, pairing aCompCor with spike regression resulted in the lowest correlations (*i*.*e*., best performance) between distance and QC-FC whether performed together with 6HMP, 24HMP, or GSR. Similarly, 24HMP with 2phys and spike regression also resulted in a low QF-FC distance dependence. Like QC-FC, the three worst pipelines all included data-driven methods (ICA-AROMA with 2phys; ICA-AROMA with 2phys and GSR; and 24HMP with aCompCor50, and GSR). Likewise, scrubbing (with 24HMP, 2phys, with or without GSR) resulted in poor performance under this exclusion regime. Overall, the inclusion of GSR worsened pipeline performance across the board. Under the (*ii*) lenient regime, the combination of aCompCor with spike regression, whether with 6 or 24HMP, resulted in very low correlations (*i*.*e*., good performance), only surpassed by the combination of ICA-AROMA with 2phys and 24HMP with 2phys and spike regression. The addition of GSR also worsened performance across all pipelines under this regime. aCompCor50 (with GSR) was the worst performer, followed by scrubbing paired with 24HMP, 2phys, and GSR, and 24HMP paired with 2phys and GSR. Finally, under the (*iii*) stringent regime, the combination of aCompCor and spike regression, whether with 6 or 24HMP, resulted in the lowest correlations (*i*.*e*., best performance). The addition of GSR generally resulted in unchanged or worse performance, with 24HMP with aCompCor50 and GSR resulting in the most significant absolute correlation.

#### 3.2.3 tDOF-Loss

As expected, the stringent regime resulted in the lowest average loss of tDOF (since the high-movement subjects were excluded), albeit at the detriment of group degrees of freedom – given the large loss of sample size. Scrubbing resulted in the most significant tDOF-loss across all regimes (**Fig. 4**).

**Fig 4:**
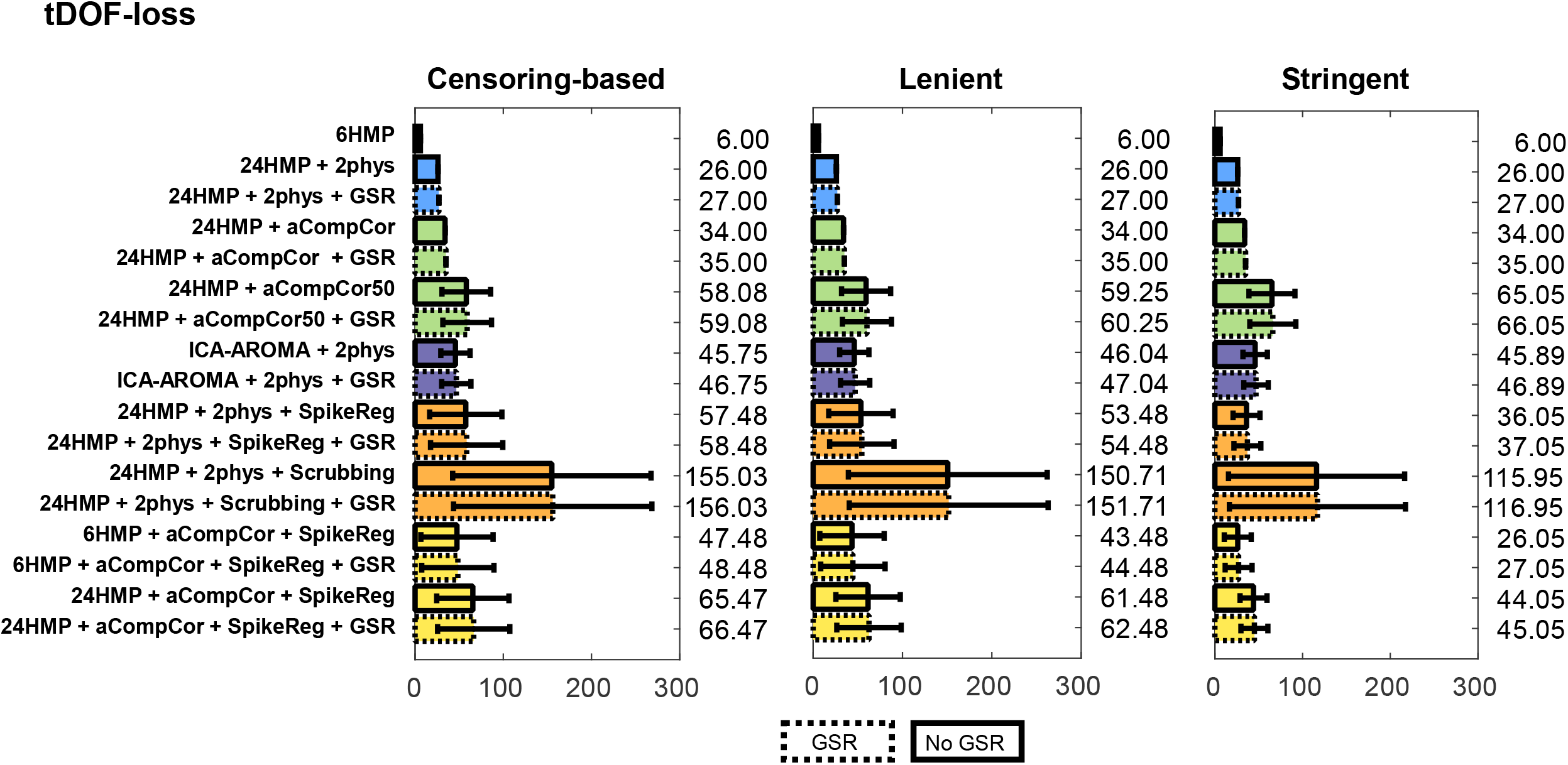
Temporal degrees of freedom loss (tDOF-loss) under the three regimes of participant exclusion. Results are presented as mean ± standard deviation. Ideally, good denoising pipelines should use fewer regressors in the model, losing fewer degrees of freedom and resulting in values closer to 0.

## 4. DISCUSSION

In this study, we report the first large-scale evaluation of denoising pipelines aimed at reducing noise-related FC in a clinical population known to be highly susceptible to in-scanner motion (Hannawi et al., 2016; Monti et al., 2015) and to present considerable anatomical abnormalities. Overall, we report three main findings.

First, one of the most critical aspects of successful denoising is selecting which subjects to retain for further analysis (Satterthwaite et al., 2013; Satterthwaite et al., 2012; Van Dijk et al., 2012). In this high-motion cohort, a stringent selection obviously resulted in equal or better performance in QC metrics across virtually all pipelines. In other words, once high-motion participants are removed from the sample, the choice of denoising pipeline becomes secondary (with the sole exception of the 6HMP approach). Nonetheless, while the quality of the data used for analysis benefits significantly from this approach, it is very costly in terms of data loss (37% in our sample). Consequently, it decreases the degrees of freedom for statistical inference across groups (such as performing group comparison between patients and volunteers or correlation analysis between behavioral scores on a test of interest and FC metrics).

Second, different denoising approaches exhibit very distinct abilities to mitigate the negative effects of noise on FC (Parkes et al., 2018). Pipelines combining spike regression with 2phys and its extension, aCompCor, tend to be the best performers across exclusion regimes. Somewhat unexpectedly, pipelines using scrubbing, ICA-AROMA, and aCompCor50 performed comparatively worse. Overall, pipelines using scrubbing were generally either comparable or worse than the other ones, in addition to the cost of two to three times greater loss of tDOF—up to 50% of the available data per subject—thus hampering the quality of the FC estimates. Pipelines containing data-driven techniques (*i*.*e*., ICA-AROMA and aCompCor50), which can be very effective at removing noise-related artifacts (Pruim et al., 2015), were instead among the worst performers under most regimes. While it is hard to pinpoint the source of their poor performance, we could speculate that the structural features of our images pose too great an obstacle to be addressed by these denoising approaches. aCompCor50 and ICA-AROMA, for example, rely on the accurate segmentation of brain tissues to identify noise components. TBI patients constitute a very heterogeneous sample from which segmenting the brain into different tissues might be challenging, probably affecting the performance of denoising strategies that depend on this step. Indeed, ICA-AROMA at times could not find any “signal” components in some patients (*i*.*e*., all components were classified as noise, **Suppl. Fig1**), stressing that these methods should be used with care when dealing with datasets containing pathological brains (Heine et al., 2012).

Third, we find the addition of GSR, a controversial step in fMRI data preprocessing, to give mixed results. On the one hand, it did improve the QC-FC metric under the lenient and stringent regimes (albeit only very marginally in the latter). On the other hand, it worsened the distance-dependent QC metric for virtually all pipelines, under all regimes—consistent with prior reports (Ciric et al., 2017).

Given these results, we offer three recommendations. First, where possible, use a stringent exclusion regime. This approach essentially reduces the analyzed sample to low-motion subjects, thus ensuring that systematic spurious correlations do not affect FC estimates. While the data loss can be sizeable (37% in our sample), this approach leads to the most significant mitigation of the negative effects of noise on FC. In addition, this strategy also gives the researcher freedom to choose among almost any pipeline, according to which procedure is best for the study’s goals. However, this approach has the potential for biased data loss. For example, some patients might be more motion-prone, resulting in greater exclusion rates and thus hampering group analyses.

Second, when choosing between pipelines, we find combinations of 2phys, spike regression and aCompCor to perform best in general. Scrubbing, in turn, performed relatively poorly under most circumstances and led to a high loss in tDOF. Likewise, ICA-AROMA—which has been shown to perform very well in healthy volunteers (Pruim et al., 2015)—underperformed many pipelines in our clinical sample. While the reason why this approach was not very effective remains unsolved, we speculate that the segmentation of tissue compartments can be very problematic in the presence of significant brain shape deformation (*e*.*g*., due to primary impact damage, ventricular enlargement, among others). Finally, given the mixed results and the controversial nature of this step, we do not recommend using GSR.

There are several limitations to the current work that should be acknowledged. First, our results are limited to the combination of approaches we chose for each pipeline. While some pipelines outperformed others, we should bear in mind that different combinations could yield divergent results (*e*.*g*., adding quadratic and derivative terms of physiological or global signal). Likewise, our results reflect the performance of pipelines for our particular image acquisition parameters. Testing these pipelines in images with shorter or longer TRs and other parameters should be addressed in future work. Second, the participants excluded from censoring pipelines (*i*.*e*., participants with < 4 min of data based on spike regression or scrubbing) were also excluded from the other pipelines. While we thought it was crucial to compare pipelines maintaining the number of subjects constant across them, it also precluded us from evaluating how non-censoring pipelines would perform without this criterion. Future work should focus on assessing, for example, how data-driven approaches perform when including these participants. Lastly, our QC measures focused on a specific way of calculating FC (*i*.*e*., a model-based method using Pearson’s correlation between ROIs time series). We recognize that other metrics of FC (see (Li et al., 2009)) could result in different findings.

Taken together, our findings stress the heterogeneous performance of denoising pipelines, emphasizing that different strategies may be appropriate in the context of specific goals, according to the question and study design. Researchers should be familiar with their samples regarding head movement profile and clinical features and be aware of each approach’s strengths and weaknesses to find the pipeline that best matches their goals.

## 5. AUTHOR CONTRIBUTIONS

The authors confirm contribution to the paper as follows: MW, RFC and MMM conceived and designed the study; the Epilepsy Bioinformatics Study for Anti-epileptogenic Therapy clinical trial (EpiBioS4Rx) research group (**Suppl. Table 4**) collected the data; MW, RFC, BMC, JSC, ESL, PMV and MMM contributed with analysis tools; MW and RFC performed the analysis; MW, RFC, BMC, JSC, ESL, PMV and MMM wrote the paper.

## FIGURE LEGENDS

**Suppl. Fig1:** Box plot showing the percentage of components classified as noise by ICA-AROMA for each patient. When 100, it means that all components were classified as noise by the classifier.

**Suppl. Table 1:** Detailed T1 image acquisition parameters:

| Subj | Mag Field | Software | Model | Matrix |  | Voxel Size |  | Slice Thickness | Slices | TR (ms) | TE (ms) | Flip Angle |
| --- | --- | --- | --- | --- | --- | --- | --- | --- | --- | --- | --- | --- |
| 1 | 3T | SIEMENS | Skyra | 256 | 256 | 1 | 1 | 1 | 256 | 2300 | 2.26 | 8 |
| 2 | 3T | SIEMENS | Skyra | 256 | 256 | 1 | 1 | 1 | 256 | 2300 | 2.26 | 8 |
| 3 | 3T | SIEMENS | Skyra | 256 | 256 | 1 | 1 | 1 | 256 | 2300 | 2.26 | 8 |
| 4 | 3T | SIEMENS | Skyra | 256 | 256 | 1 | 1 | 1 | 256 | 2300 | 2.26 | 8 |
| 5 | 3T | SIEMENS | Skyra | 256 | 256 | 1 | 1 | 1 | 256 | 2300 | 2.26 | 8 |
| 6 | 3T | SIEMENS | Skyra | 256 | 256 | 1 | 1 | 1 | 256 | 2300 | 2.26 | 8 |
| 7 | 3T | SIEMENS | Skyra | 256 | 256 | 1 | 1 | 1 | 256 | 2300 | 2.26 | 8 |
| 8 | 3T | SIEMENS | Skyra | 256 | 256 | 1 | 1 | 1 | 256 | 2300 | 2.26 | 8 |
| 9 | 3T | SIEMENS | Skyra | 256 | 256 | 1 | 1 | 1 | 256 | 2300 | 2.26 | 8 |
| 10 | 3T | SIEMENS | Skyra | 256 | 256 | 1 | 1 | 1 | 256 | 2300 | 2.26 | 8 |
| 11 | 3T | SIEMENS | Skyra | 256 | 256 | 1 | 1 | 1 | 256 | 2300 | 2.26 | 8 |
| 12 | 3T | Philips | Ingenia | 256 | 256 | 1 | 1 | 1 | 256 | 8.196 | 3.753 | 8 |
| 13 | 3T | Philips | Ingenia | 256 | 256 | 1 | 1 | 1 | 256 | 8.247 | 3.776 | 8 |
| 14 | 3T | Philips | Ingenia | 256 | 256 | 1 | 1 | 1 | 256 | 8.142 | 3.721 | 8 |
| 15 | 3T | Philips | Ingenia | 280 | 220 | 0.49 | 0.49 | 0.9 | 512 | 9.01 | 4.124 | 8 |
| 16 | 3T | Philips | Ingenia | 256 | 256 | 1 | 1 | 1 | 256 | 8.233 | 3.771 | 8 |
| 17 | 3T | Philips | Ingenia | 256 | 256 | 1 | 1 | 1 | 256 | 8.397 | 3.86 | 8 |
| 18 | 3T | GE | Signa | 256 | 256 | 1 | 1 | 1 | 256 | 8.856 | 3.488 | 15 |
| 19 | 3T | GE | Signa | 256 | 256 | 1 | 1 | 1 | 256 | 9.088 | 3.624 | 15 |
| 20 | 3T | GE | Signa | 256 | 256 | 1 | 1 | 1 | 256 | 8.824 | 3.488 | 15 |
| 21 | 3T | GE | Signa | 256 | 256 | 1 | 1 | 1 | 256 | 8.86 | 3.488 | 15 |
| 22 | 3T | GE | Signa | 256 | 256 | 1 | 1 | 1 | 256 | 8.868 | 3.488 | 15 |
| 23 | 3T | GE | Signa | 256 | 256 | 1 | 1 | 1 | 256 | 9.148 | 3.624 | 15 |
| 24 | 3T | GE | Signa | 256 | 256 | 1 | 1 | 1 | 256 | 11.28 | 4.848 | 20 |
| 25 | 3T | GE | Signa | 256 | 256 | 1 | 1 | 1 | 256 | 8.836 | 3.488 | 15 |
| 26 | 3T | GE | Signa | 256 | 256 | 1 | 1 | 1 | 256 | 9.116 | 3.524 | 15 |
| 27 | 3T | GE | Signa | 256 | 256 | 1 | 1 | 1 | 256 | 9.128 | 3.624 | 15 |
| 28 | 3T | GE | Signa | 256 | 256 | 1 | 1 | 1 | 256 | 9.136 | 3.624 | 15 |
| 29 | 3T | GE | Signa | 256 | 256 | 1 | 1 | 1 | 256 | 9.148 | 3.624 | 15 |
| 30 | 3T | GE | Signa | 256 | 256 | 1 | 1 | 1 | 256 | 9.076 | 3.624 | 15 |
| 31 | 3T | GE | Signa | 256 | 256 | 1 | 1 | 1 | 256 | 9.128 | 3.624 | 15 |
| 32 | 3T | GE | Signa | 256 | 256 | 1 | 1 | 1 | 256 | 9.148 | 3.616 | 15 |
| 33 | 3T | GE | Signa | 256 | 256 | 1 | 1 | 1 | 256 | 9.12 | 3.624 | 15 |
| 34 | 3T | GE | Signa | 256 | 256 | 1 | 1 | 1 | 256 | 9.096 | 3.624 | 15 |
| 35 | 3T | GE | Signa | 256 | 256 | 1 | 1 | 1 | 256 | 9.048 | 3.616 | 15 |
| 36 | 3T | GE | Signa | 256 | 256 | 1 | 1 | 1 | 256 | 9.096 | 3.616 | 15 |
| 37 | 3T | GE | Signa | 256 | 256 | 1 | 1 | 1 | 256 | 9.096 | 3.616 | 15 |
| 38 | 3T | GE | Signa | 256 | 256 | 1 | 1 | 1 | 256 | 9.148 | 3.616 | 15 |
| 39 | 3T | SIEMENS | TrioTim | 256 | 256 | 1 | 1 | 1 | 160 | 1900 | 3.43 | 9 |
| 40 | 3T | SIEMENS | TrioTim | 256 | 256 | 1 | 1 | 1 | 160 | 1900 | 3.43 | 9 |
| 41 | 3T | SIEMENS | TrioTim | 256 | 256 | 1 | 1 | 1 | 160 | 1900 | 3.43 | 9 |
| 42 | 3T | SIEMENS | TrioTim | 256 | 256 | 1 | 1 | 1 | 159 | 1900 | 3.43 | 9 |
| 43 | 3T | SIEMENS | TrioTim | 256 | 256 | 1 | 1 | 1 | 160 | 1900 | 3.43 | 9 |
| 44 | 3T | SIEMENS | TrioTim | 256 | 256 | 1 | 1 | 1 | 154 | 1900 | 3.43 | 9 |
| 45 | 3T | SIEMENS | TrioTim | 256 | 256 | 1 | 1 | 1 | 160 | 1900 | 3.43 | 9 |
| 46 | 3T | SIEMENS | TrioTim | 256 | 256 | 1 | 1 | 1 | 160 | 1900 | 3.43 | 9 |
| 47 | 3T | SIEMENS | TrioTim | 256 | 256 | 1 | 1 | 1 | 160 | 1900 | 3.43 | 9 |
| 48 | 3T | SIEMENS | TrioTim | 256 | 256 | 1 | 1 | 1 | 160 | 1900 | 3.43 | 9 |
| 49 | 3T | SIEMENS | TrioTim | 256 | 256 | 1 | 1 | 1 | 160 | 1900 | 3.43 | 9 |
| 50 | 3T | SIEMENS | TrioTim | 256 | 256 | 1 | 1 | 1 | 160 | 1900 | 3.43 | 9 |
| 51 | 3T | SIEMENS | TrioTim | 256 | 256 | 1 | 1 | 1 | 160 | 1900 | 3.43 | 9 |
| 52 | 3T | SIEMENS | TrioTim | 256 | 256 | 1 | 1 | 1 | 160 | 1900 | 3.43 | 9 |
| 53 | 3T | SIEMENS | TrioTim | 256 | 256 | 1 | 1 | 1 | 160 | 1900 | 3.43 | 9 |
| 54 | 3T | SIEMENS | TrioTim | 256 | 256 | 1 | 1 | 1 | 160 | 1900 | 3.43 | 9 |
| 55 | 3T | SIEMENS | TrioTim | 256 | 256 | 1 | 1 | 1 | 160 | 1900 | 3.43 | 9 |
| 56 | 3T | SIEMENS | TrioTim | 256 | 256 | 1 | 1 | 1 | 160 | 1900 | 3.43 | 9 |
| 57 | 3T | SIEMENS | TrioTim | 256 | 256 | 1 | 1 | 1 | 160 | 1900 | 3.43 | 9 |
| 58 | 3T | SIEMENS | TrioTim | 256 | 256 | 1 | 1 | 1 | 160 | 1900 | 3.43 | 9 |
| 59 | 3T | SIEMENS | TrioTim | 256 | 256 | 1 | 1 | 1 | 160 | 1900 | 3.43 | 9 |
| 60 | 3T | SIEMENS | TrioTim | 256 | 256 | 1 | 1 | 1 | 160 | 1900 | 3.43 | 9 |
| 61 | 1.5T | SIEMENS | Aera | 256 | 256 | 1 | 1 | 1 | 160 | 2000 | 3.13 | 15 |
| 62 | 1.5T | SIEMENS | Aera | 256 | 256 | 1 | 1 | 1 | 192 | 2000 | 3.13 | 15 |
| 63 | 1.5T | SIEMENS | Aera | 256 | 256 | 1 | 1 | 1 | 160 | 2000 | 3.13 | 15 |
| 64 | 1.5T | SIEMENS | Aera | 256 | 256 | 1 | 1 | 1 | 144 | 2000 | 3.13 | 15 |
| 65 | 1.5T | SIEMENS | Aera | 256 | 256 | 1 | 1 | 1 | 160 | 2000 | 3.13 | 15 |
| 66 | 1.5T | SIEMENS | Aera | 256 | 256 | 1 | 1 | 1 | 128 | 2000 | 3.13 | 15 |
| 67 | 1.5T | SIEMENS | Aera | 256 | 256 | 1 | 1 | 1 | 144 | 2000 | 3.13 | 15 |
| 68 | 1.5T | SIEMENS | Aera | 256 | 256 | 1 | 1 | 1 | 176 | 2000 | 3.13 | 15 |
| 69 | 1.5T | SIEMENS | Aera | 256 | 256 | 1 | 1 | 1 | 176 | 2000 | 3.13 | 15 |
| 70 | 1.5T | SIEMENS | Aera | 256 | 256 | 1 | 1 | 1 | 128 | 2000 | 3.13 | 15 |
| 71 | 1.5T | SIEMENS | Aera | 256 | 256 | 1 | 1 | 1 | 160 | 2000 | 3.13 | 15 |
| 72 | 1.5T | SIEMENS | Aera | 256 | 256 | 1 | 1 | 1 | 176 | 2000 | 3.13 | 15 |
| 73 | 1.5T | SIEMENS | Aera | 256 | 256 | 1 | 1 | 1 | 128 | 2000 | 3.13 | 15 |
| 74 | 1.5T | SIEMENS | Aera | 256 | 256 | 1 | 1 | 1 | 192 | 2000 | 3.13 | 15 |
| 75 | 1.5T | SIEMENS | Aera | 256 | 256 | 1 | 1 | 1 | 144 | 2000 | 3.13 | 15 |
| 76 | 1.5T | SIEMENS | Aera | 256 | 256 | 1 | 1 | 1 | 144 | 2000 | 3.13 | 15 |
| 77 | 1.5T | SIEMENS | Aera | 256 | 256 | 1 | 1 | 1 | 144 | 2000 | 3.13 | 15 |
| 78 | 3T | SIEMENS | Verio | 256 | 256 | 1 | 1 | 1 | 160 | 1900 | 2.93 | 9 |
| 79 | 3T | SIEMENS | Verio | 256 | 256 | 1 | 1 | 1 | 160 | 1900 | 2.93 | 9 |
| 80 | 3T | SIEMENS | Verio | 256 | 256 | 1 | 1 | 1 | 176 | 1900 | 2.93 | 9 |
| 81 | 3T | SIEMENS | Verio | 256 | 256 | 1 | 1 | 1 | 160 | 1900 | 2.93 | 9 |
| 82 | 3T | SIEMENS | Verio | 256 | 256 | 1 | 1 | 1 | 176 | 1900 | 2.93 | 9 |
| 83 | 3T | SIEMENS | Skyra | 256 | 256 | 1 | 1 | 1 | 256 | 2300 | 2.26 | 8 |
| 84 | 3T | SIEMENS | Skyra | 256 | 256 | 1 | 1 | 1 | 256 | 2300 | 2.26 | 8 |
| 85 | 3T | SIEMENS | Skyra | 256 | 256 | 1 | 1 | 1 | 256 | 2300 | 2.26 | 8 |
| 86 | 3T | SIEMENS | Skyra | 256 | 256 | 1 | 1 | 1 | 256 | 2300 | 2.26 | 8 |
| 87 | 3T | SIEMENS | Skyra | 256 | 256 | 1 | 1 | 1 | 256 | 2300 | 2.26 | 8 |
| 88 | 3T | SIEMENS | Skyra | 256 | 256 | 1 | 1 | 1 | 256 | 2300 | 2.26 | 8 |

**Suppl. Table 2:**
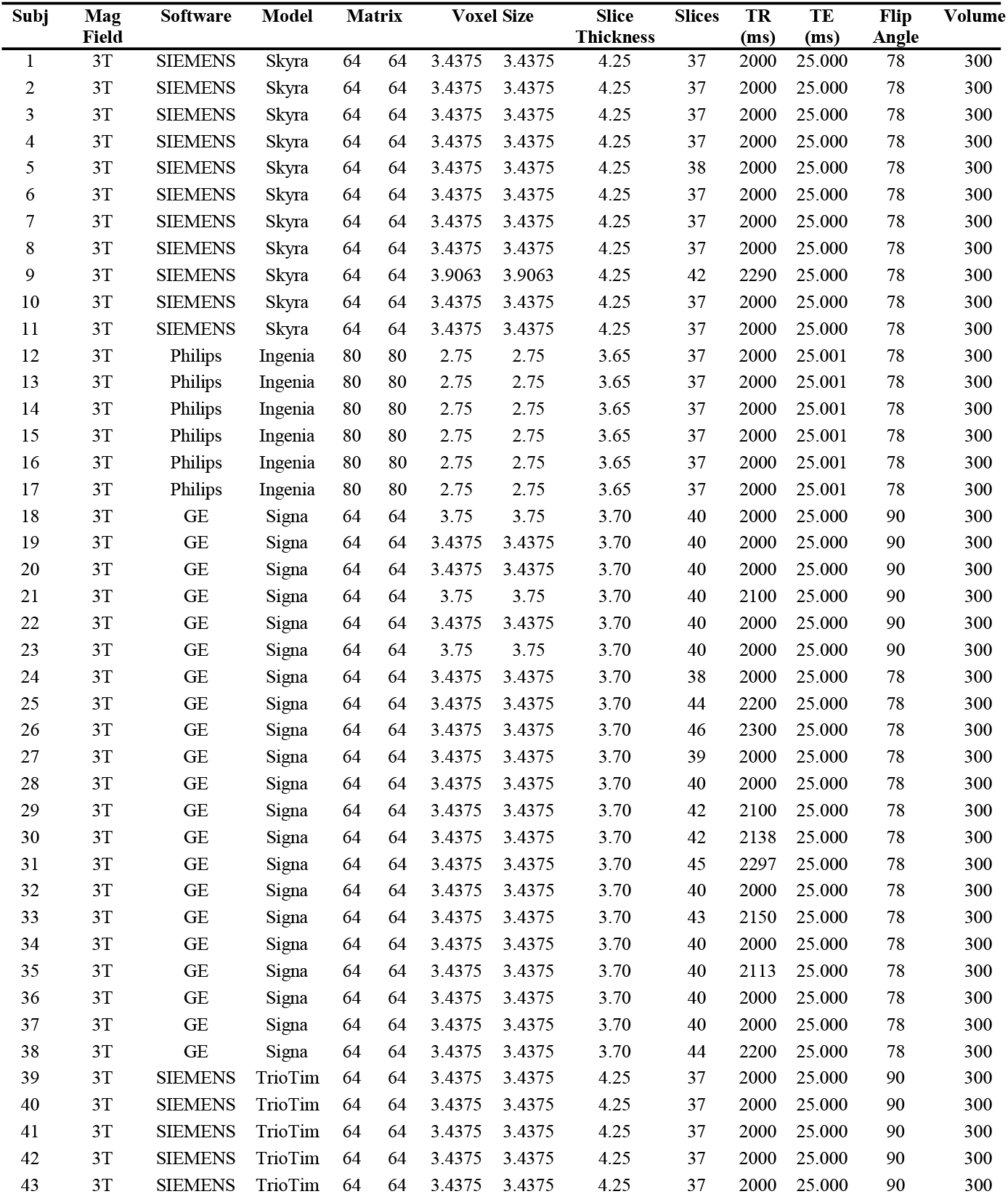
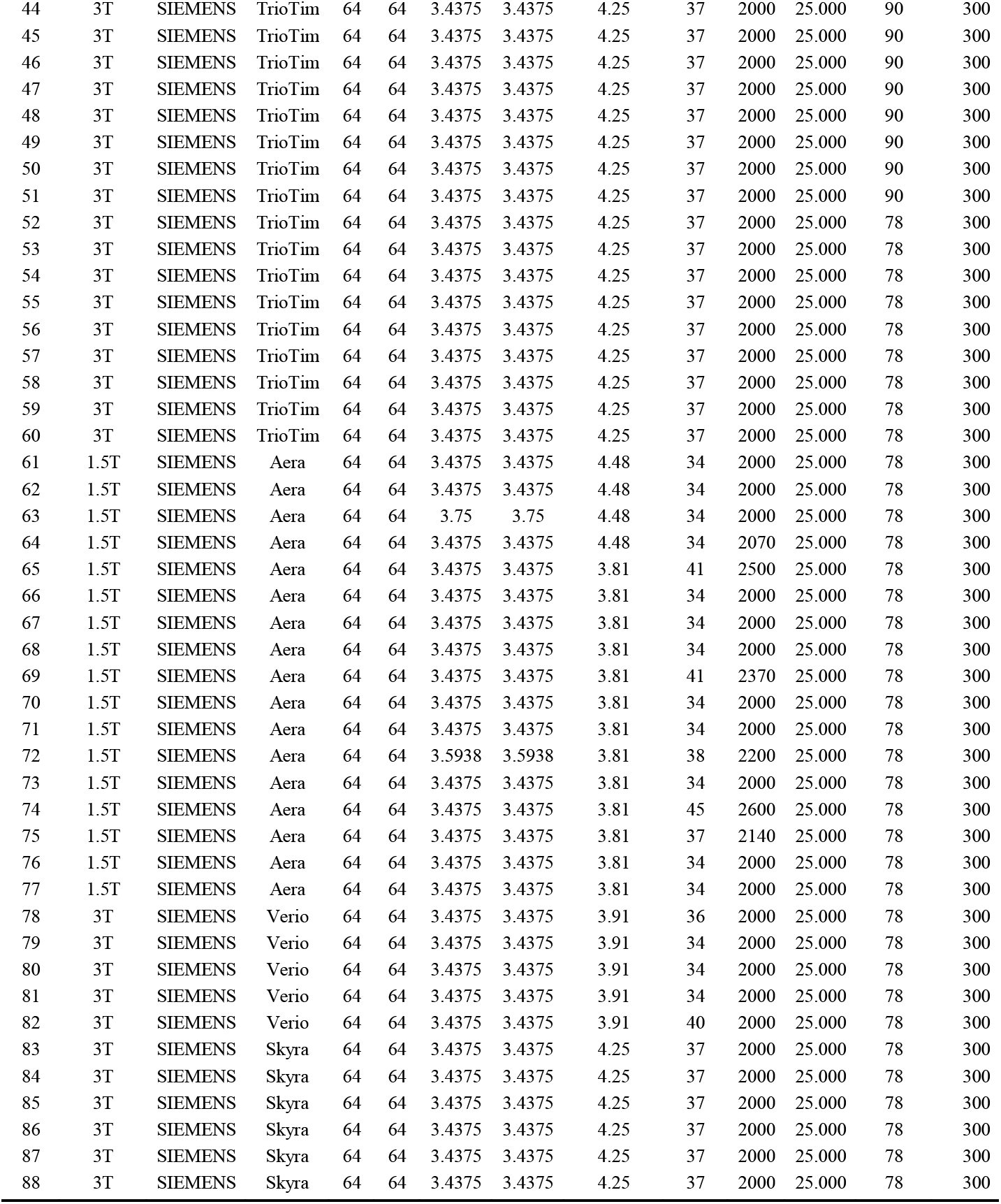
Detailed T2*-weighted echo planar images acquisition parameters:

**Suppl. Table 3:**
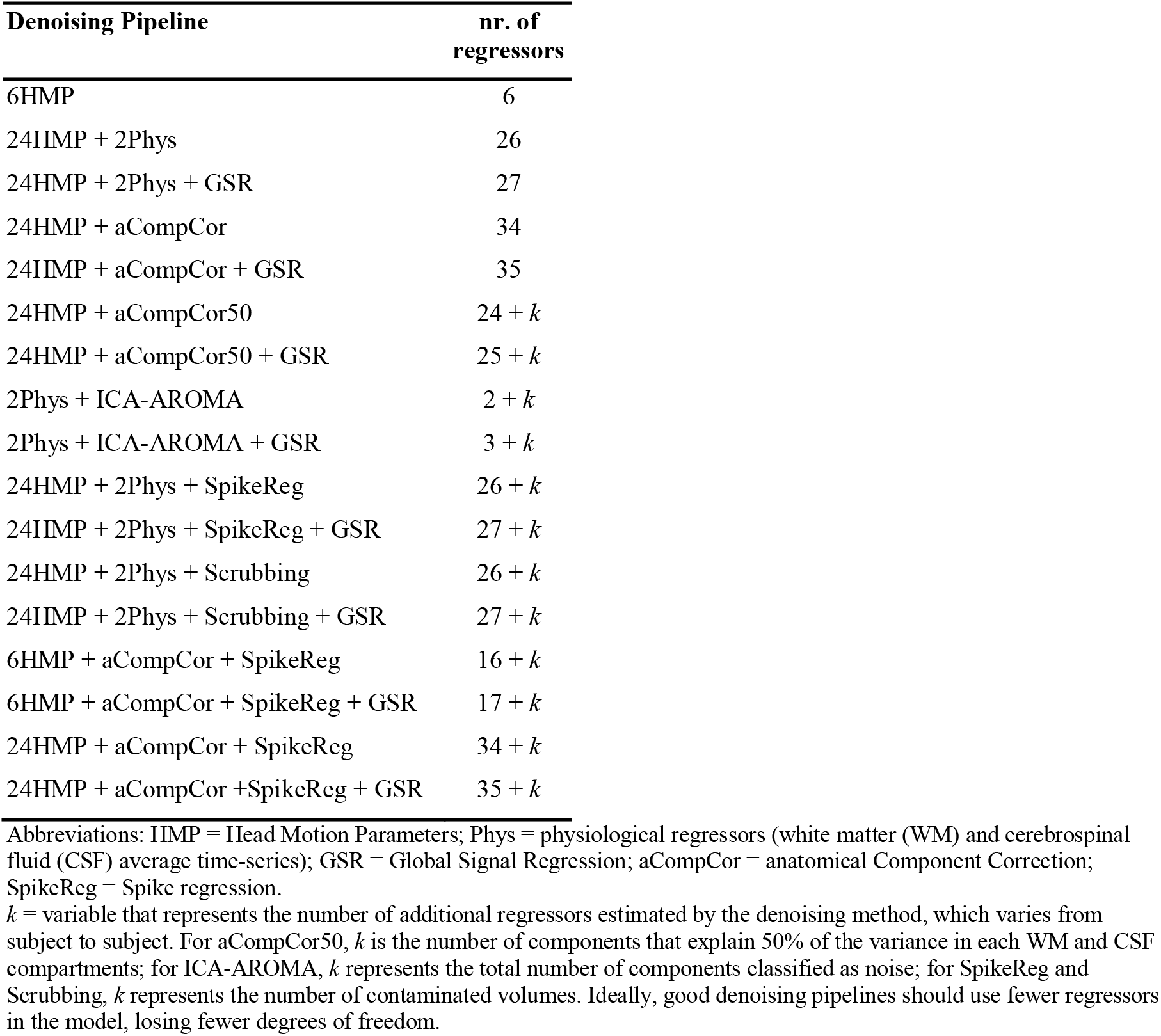
Denoising pipelines and the total number of regressors used in each of them.

**Suppl. Table 4:** EpiBioS4Rx’s Principal Investigators and affiliated institutions.

| Principal Investigator/Author | Affiliated Institutions |
| --- | --- |
| Alaa Kamnaksh | Uniformed Services University, United States |
| Alicia Au | University of Pittsburgh, United States |
| Andrew Morokoff | Royal Melbourne Hospital, United States |
| Arthur Toga | University of Southern California, United States |
| Ben Ellingson | David Geffen School of Medicine at UCLA, United States |
| Brandon Foreman | University of Cincinnati, United States |
| Courtney L. Robertson | Johns Hopkins University, United States |
| Courtney Real | David Geffen School of Medicine at UCLA, United States |
| David McArthur | David Geffen School of Medicine at UCLA, United States |
| Denes Agoston | Uniformed Services University, United States |
| Elisa Yam | David Geffen School of Medicine at UCLA, United States |
| Emily J. Gilmore | Yale University, United States |
| Eric Rosenthal | Harvard University/Massachusetts General Hospital, United States |
| Jan Claassen | Columbia University, United States |
| Lara Zimmermann | University of California, Davis, United States |
| Lawrence Hirsch | Yale University, United States |
| Manuel Buitrago Blanco | David Geffen School of Medicine at UCLA, United States |
| Michael J. Bell | Children's National Health System, United States |
| Ramon Diaz-Arrastia | University of Pennsylvania, United States |
| Richard Staba | David Geffen School of Medicine at UCLA, United States |
| Susana Martinez | David Geffen School of Medicine at UCLA, United States |
| Terrence O'Brien | Royal Melbourne Hospital, United States |
| Dominique Duncan | University of Southern California, United States |
| Frederick A. Willyerd | Phoenix Children's Hospital, United States |
| Jerome Engel Jr. | David Geffen School of Medicine at UCLA, United States |
| Kristine O'Phelan | University of Miami, United States |
| Martin Hunn | The Alfred Hospital Melbourne, Australia |
| Nicholas Abend | University of Pennsylvania, United States |
| Thomas P. Bleck | Rush University, United States |
| Vikesh Shrestha | David Geffen School of Medicine at UCLA, United States |

