## Supplementary figures and images for "Evaluating denoising strategies in resting-state fMRI in traumatic brain injury (EpiBioS4Rx)"

### Supplemental Figure 1

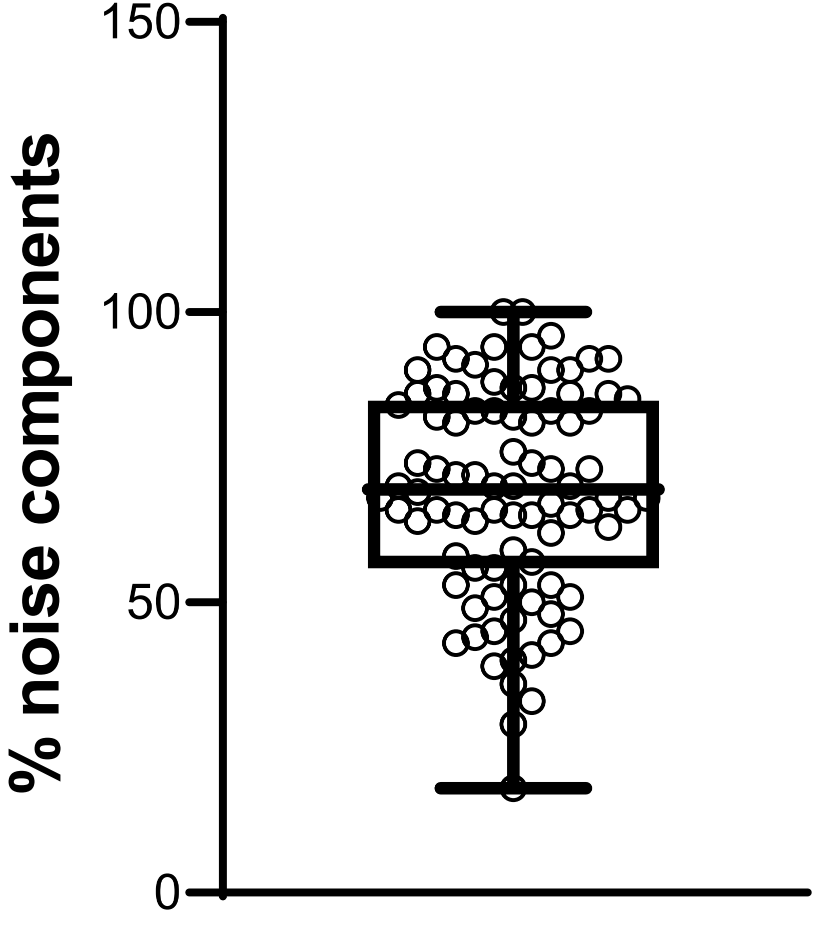
